# Physical contact reveals a hidden layer of cortical architecture

**DOI:** 10.64898/2026.05.08.723866

**Authors:** Jordan K. Matelsky, Hannah Martinez, Michael S. Robinette, Katharine Merfeld, Daniel Xenes, Cara J. Cavanaugh, Sarah E. Emerson, Dhananjay Bhaskar, Benjamin Clark, Caitlyn Bishop, Konrad P. Körding, Daniel Colón-Ramos, Patricia Rivlin, Cody J. Smith, Brock Wester

## Abstract

Neurons interact at synapses, but they also communicate through physical contact and proximity, including diffusion, glia-mediated interactions, and ephaptic coupling. Standard connectomes map synapses, but cannot capture the full set of cell-cell contacts that can support these pathways. Here we extract contactomes from two large mouse visual cortex volumes at nanoscale resolution and quantify every cell-cell contact, the shared surface area of each contact, and the relationship between contact and synaptic connectivity. We find that contactomes are 5 − 10× denser than synaptic graphs, revealing that neurons physically contact a much larger set of potential neighbors than they synaptically connect to. We further find that most nearby potential neighbors are already in physical contact, indicating that local structural change would add few new candidate synaptic partners. Finally, we find that astrocytes form a single large syncytium-like network that spans the tissue and directly contacts nearly all neurons, and that glial processes lie within a micron or two of almost every synapse, indicating that synapses reside within a pervasive glia-shaped microenvironment. Together, these results show that physical contact forms a distinct layer of brain architecture that extends far beyond the synaptic connectome.

## Introduction

Evolution has balanced metabolic and computational pressures to build brains that are highly connected, tightly packed, and yet energy-efficient.^1–3^ The function of a cell depends on its relationships to surrounding cells, and synapses are only one abstraction of those relationships. The nervous system takes advantage of diverse forms of intercellular communication in addition to chemical synapses — such as gap junctions, hormonal exchange, neuropeptide “wireless” communication, and ephaptic coupling — and these systems all play a role in cellular communication and computational organization.^2,4–12^ But synaptic network connectomes cannot capture this broader field. To understand these relationships at scale, we need exhaustive maps of cellular apposition.

Networks of cellular apposition, unlike the synaptic connectome, can include *all* cells in the nervous system, spanning neurons as well as glia, pericytes, and endothelium, making it possible to analyze neuron-glia organization and tissue-scale non-neuronal structure within the same relational framework. Despite the importance of such systems, exhaustive contactome maps have only been available for small nervous systems like *C. elegans*^2,13–17^ and for specialized subcellular structures within larger nervous systems.^18^ Those more localized or specialized studies established that contact structure can reveal organizational features that are not visible in the synaptic graph alone, but comparable exhaustive whole-cell contactome analyses have not previously been extended to mammalian cortex at this scale and complexity. Together with a companion paper^19^, we introduce a computationally simpler way to derive contactome graphs from even petabyte-scale datasets using a chunked geometric operation on piecewise connected-components. This representation includes cell surface area measurements between all cells in the volume. This makes it possible to study contact structure in large cortical electron microscopy (EM) volumes, including, to our knowledge, the first saturated, whole-cell contactome analysis of mammalian cortex at cubic-millimeter scale. It also enables quantitative analysis of neuron-neuron apposition within one unified whole-cell contact graph, including pairwise shared surface area, alongside the first analyses at this scale of astrocyte network organization and near-synapse glial context.

We show three main results: (1) Cell-cell contactomes are five to ten times denser than synaptic graphs, exposing many physically available neuron-neuron relationships that are not realized as synapses (**Fig. 1A,C,D**). (2) Astrocytes form a large syncytium-like contact network that spans the tissue and places nearly all neurons in direct contact with this glial network. This organization is consistent with tissue-scale astrocytic coupling (**Fig. 1B**). (3) Nearly every synapse (99%) lies within 2 µm of at least one glial process, indicating that most synapses reside within a local microenvironment shaped by glia (**Fig. 1E**).

**Figure 1.**
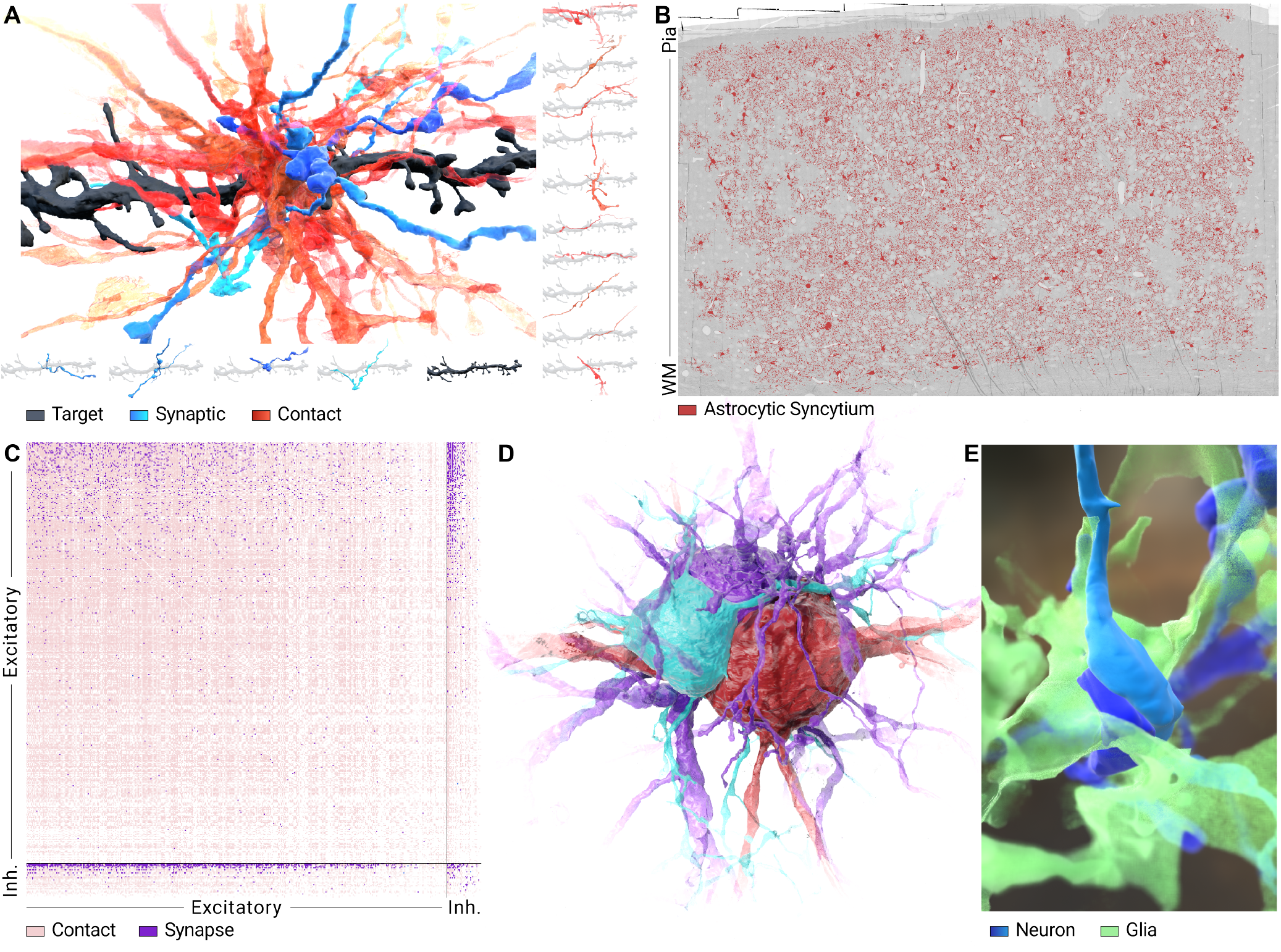
The contactome reveals the rich structure of the neocortex architecture. **A. Exemplar dendritic spine with both its synaptic partners and nearby contact neighbors, each shown in isolation**. A single dendritic spine from a target process (black) has four synaptic partners (blue, shown below) but more than fifteen contact neighbors (red, subsampled in exploded view right). **B. Astrocytes form a syncytium-like network that spans the cortical volume**. This dominant astrocyte network directly contacts most neurons in the volume. The displayed tissue comprises the complete 1mm volume. **C. Comparison of synaptic and contact adjacency matrices**. The traditional synaptic connectome adjacency matrix of the MICrONS Layer 2/3 volume is shown in purple. Contact adjacency (red) is substantially denser, and *also* includes nearly all synaptic edges. **D. Three neural somas make intimate contact**. A large fraction of total surface area of these three somas is shared, though this interaction is completely invisible in the synaptic connectome. **E. Synapses near universally (99%) have glia within a distance or two microns**. This cutaway shows a glial process near a synapse, illustrating the local microenvironment discussed more in *Results*. High resolution images and links to interactive versions are available in the *Supplemental Materials*.

## Results

### Contact exposes far more neighbors than synapses alone

We can think of synapses as a subset of the full field of possible interactions between cells. Because many forms of intercellular communication require physical contact, the contactome should expose a much larger pool of neuron-neuron neighbors than synapses do. Reviewing the MICrONS Layer 2/3^20^ and the MICrONS 1 *mm*^321^ mouse visual cortex volumes, we found that contact density exceeded synapse density by 11.74× in the smaller volume and 3.853× in the larger volume, consistent with previous estimates (**Fig. 1A**).^22^ On average, neurons encountered thousands to tens of thousands of unique other cells via contact, and only made synapses with hundreds to thousands of unique other neurons (**Fig. 2**). Subtype-level full-graph contact degree and total contact-area statistics are summarized in **Table 1**.

**Table 1.** Contact degree and total contact-area statistics per cell type in the full mouse visual cortex contactome. Rows are grouped by cell type, and degree and contact area are measured against all contact partners in the full graph, including untyped cells. *\* Pericyte values should be interpreted cautiously because pericyte segmentation is lower-quality than that of other cell types in this volume*.

| Subtype | n | Mean degree | Median degree | Mean area ($\mu\text{m}^2$) | Median area ($\mu\text{m}^2$) |
| --- | --- | --- | --- | --- | --- |
| 23P | 19,735 | 18,906.6 | 17,730 | $4.226 \times 10^8$ | $3.241 \times 10^8$ |
| 4P | 14,777 | 8,683.7 | 5,907 | $1.891 \times 10^8$ | $1.037 \times 10^8$ |
| 5P-ET | 2,215 | 19,456.2 | 17,521 | $4.732 \times 10^8$ | $3.668 \times 10^8$ |
| 5P-IT | 7,949 | 10,852.2 | 9,061 | $2.303 \times 10^8$ | $1.575 \times 10^8$ |
| 5P-NP | 970 | 7,684.5 | 5,859.5 | $1.573 \times 10^8$ | $1.132 \times 10^8$ |
| 6P-CT | 6,815 | 8,535.0 | 7,683 | $2.034 \times 10^8$ | $1.463 \times 10^8$ |
| 6P-IT | 11,734 | 9,799.4 | 8,704 | $2.405 \times 10^8$ | $1.718 \times 10^8$ |
| BC | 3,288 | 15,111.2 | 9,064.5 | $3.346 \times 10^8$ | $1.671 \times 10^8$ |
| BPC | 1,721 | 6,748.5 | 5,520 | $1.403 \times 10^8$ | $1.010 \times 10^8$ |
| MC | 2,446 | 11,675.0 | 7,467.5 | $2.511 \times 10^8$ | $1.330 \times 10^8$ |
| NGC | 508 | 11,864.6 | 9,261 | $2.353 \times 10^8$ | $1.503 \times 10^8$ |
| astrocyte | 7,838 | 15,037.4 | 7,755 | $8.723 \times 10^8$ | $2.219 \times 10^8$ |
| microglia | 2,711 | 6,458.5 | 2,503 | $2.050 \times 10^8$ | $5.223 \times 10^7$ |
| oligo | 7,029 | 4,173.6 | 2,336 | $8.166 \times 10^7$ | $3.196 \times 10^7$ |
| OPC | 1,638 | 7,534.8 | 5,318.5 | $1.753 \times 10^8$ | $9.131 \times 10^7$ |
| pericyte* | 2,640 | 8,306.3 | 944.5 | $1.105 \times 10^9$ | $1.053 \times 10^8$ |

**Table 2.** Neuronal type-pair synaptic filling fractions in the Minnie mouse visual cortex volume. Each entry reports the percentage of contacting cell pairs of the indicated types that also share at least one synapse.

| Type | 23P (%) | 4P (%) | 5P-IT (%) | 5P-ET (%) | 5P-NP (%) | 6P-IT (%) | 6P-CT (%) | BC (%) | BPC (%) | MC (%) | NGC (%) |
| --- | --- | --- | --- | --- | --- | --- | --- | --- | --- | --- | --- |
| 23P | 4.33 | 4.51 | 5.09 | 7.79 | 2.47 | 4.97 | 4.13 | 27.59 | 8.07 | 23.70 | 14.72 |
| 4P | 4.51 | 5.84 | 6.77 | 9.64 | 3.12 | 5.68 | 5.30 | 25.97 | 9.49 | 22.87 | 8.71 |
| 5P-IT | 5.09 | 6.77 | 6.44 | 8.46 | 4.05 | 4.50 | 4.90 | 21.08 | 10.73 | 15.43 | 9.65 |
| 5P-ET | 7.79 | 9.64 | 8.46 | 10.09 | 6.97 | 6.81 | 7.55 | 24.79 | 12.55 | 17.06 | 12.53 |
| 5P-NP | 2.47 | 3.12 | 4.05 | 6.97 | 4.28 | 5.16 | 6.00 | 9.15 | 8.09 | 8.86 | 5.10 |
| 6P-IT | 4.97 | 5.68 | 4.50 | 6.81 | 5.16 | 5.09 | 5.29 | 16.52 | 7.09 | 9.72 | 11.53 |
| 6P-CT | 4.13 | 5.30 | 4.90 | 7.55 | 6.00 | 5.29 | 5.53 | 15.57 | 7.22 | 9.02 | 10.78 |
| BC | 27.59 | 25.97 | 21.08 | 24.79 | 9.15 | 16.52 | 15.57 | 34.44 | 21.49 | 26.19 | 20.93 |
| BPC | 8.07 | 9.49 | 10.73 | 12.55 | 8.09 | 7.09 | 7.22 | 21.49 | 19.64 | 24.25 | 22.22 |
| MC | 23.70 | 22.87 | 15.43 | 17.06 | 8.86 | 9.72 | 9.02 | 26.19 | 24.25 | 21.50 | 22.36 |
| NGC | 14.72 | 8.71 | 9.65 | 12.53 | 5.10 | 11.53 | 10.78 | 20.93 | 22.22 | 22.36 | 27.24 |

**Table 3.** Neuron coverage by the largest mouse visual cortex astrocyte syncytium component, stratified by neuron cell type. Counts report typed neurons that are directly apposed to at least one astrocyte in the largest astrocyte connected component.

| Cell type | *N* neurons | *N* touched | *N* not touched | Touched (%) |
| --- | --- | --- | --- | --- |
| 6P-IT | 11,734 | 10,693 | 1,041 | 91.13 |
| 5P-NP | 970 | 842 | 128 | 86.80 |
| 6P-CT | 6,815 | 5,881 | 934 | 86.29 |
| 23P | 19,735 | 16,746 | 2,989 | 84.85 |
| 5P-IT | 7,949 | 6,659 | 1,290 | 83.77 |
| 5P-ET | 2,215 | 1,798 | 417 | 81.17 |
| BPC | 1,721 | 1,373 | 348 | 79.78 |
| 4P | 14,777 | 11,699 | 3,078 | 79.17 |
| MC | 2,446 | 1,838 | 608 | 75.14 |
| NGC | 508 | 381 | 127 | 75.00 |
| BC | 3,288 | 2,285 | 1,003 | 69.50 |

**Figure 2.**
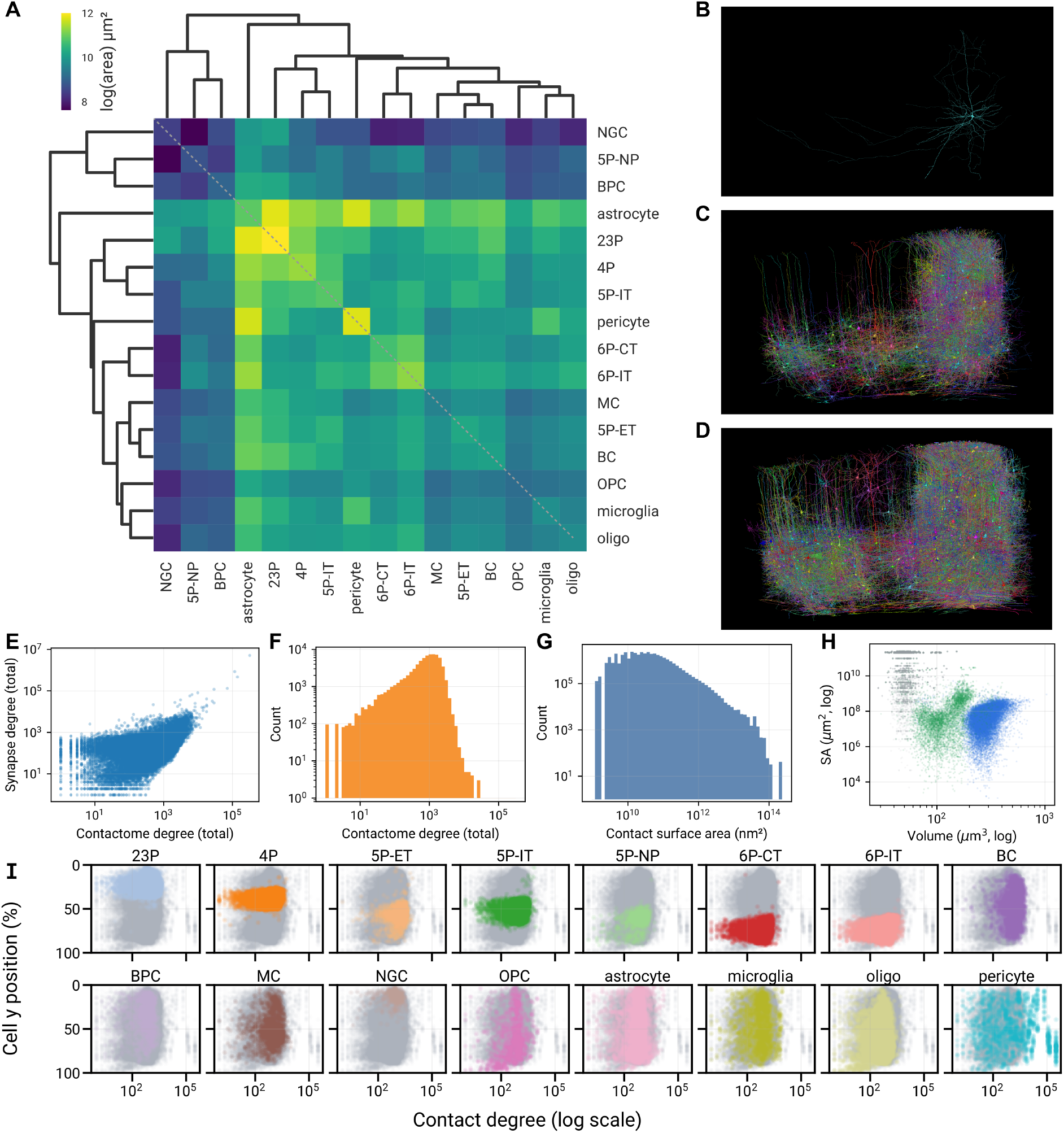
Characteristics of the cubic-millimeter mouse visual cortex contactome. **A**. Adjacency matrix of the contactome, grouped by cell type. Cell types are clustered by similarity of their contact profiles. **B**. One representative pyramidal neuron, its synaptic partners **(C)**, and its contact neighbors **(D)**. These contact neighbors span nearly the complete spatial extent of the dataset. **E**. Contact degree versus synapse degree across cells. **F**. Distribution of contact degree. **G**. Distribution of pairwise contact surface area. **H**. Cell volume versus total contact surface area. Blue points are neurons and green points are glia. **I**. Contact degree by cortical depth and cell type. Different cell types show different depth and contact degree profiles.

Despite their large difference in size, the two volumes devoted a similar order of magnitude of membrane area per tissue volume: 2.573 × 10^4^ and 4.395 × 10^4^ cm^2^/mm^3^ in the Layer 2/3 and 1 *mm*^3^ volumes, respectively. Yet only 3.41% of contacting pairs in the Layer 2/3 volume and 2.81% in the 1 *mm*^3^ volume also shared at least one synapse, indicating a quite low synaptic *filling fraction* of the contact graph in both datasets. Within the 1 *mm*^3^ volume, this filling fraction varied strongly by subtype pair, from 2.47% for 23P↔5P-NP pairs to 34.44% for BC↔BC pairs; excitatory–excitatory pairs averaged 5.27%, excitatory–inhibitory pairs 19.19%, and inhibitory–inhibitory pairs 26.74% (*Supplementary Table 2*).

### Nearby contact neighbors are already realized

Synaptogenesis is thought to continue throughout an animal’s life but is limited by the spatial contacts of two potential synaptic partners. We therefore asked whether local structural remodeling would mostly expose new potential synaptic partners or elaborate existing ones. To answer this, we defined fixed-distance halos around each neuron and captured the sets of other cells whose minimum distance fell within each halo. We then compared those nearby cells with the neighbors each neuron already contacted. We found that the median neuron made contact with 89.85% of the cells within its 1 *µ*m halo, and with 79.87% of the cells within 5 *µ*m. Even within its 10 *µ*m halo — nearly a 1000-fold increase in volume over the neuron itself — the median pyramidal neuron already contacted 75.47% of the cells nearby (**Fig. 3**). Therefore, the nearby tissue is not an untapped reservoir of neighbors, but a densely connected local neighborhood in which local remodeling is more likely to elaborate existing relationships than to create truly *new*, previously-unseen neighbors.

**Figure 3.**
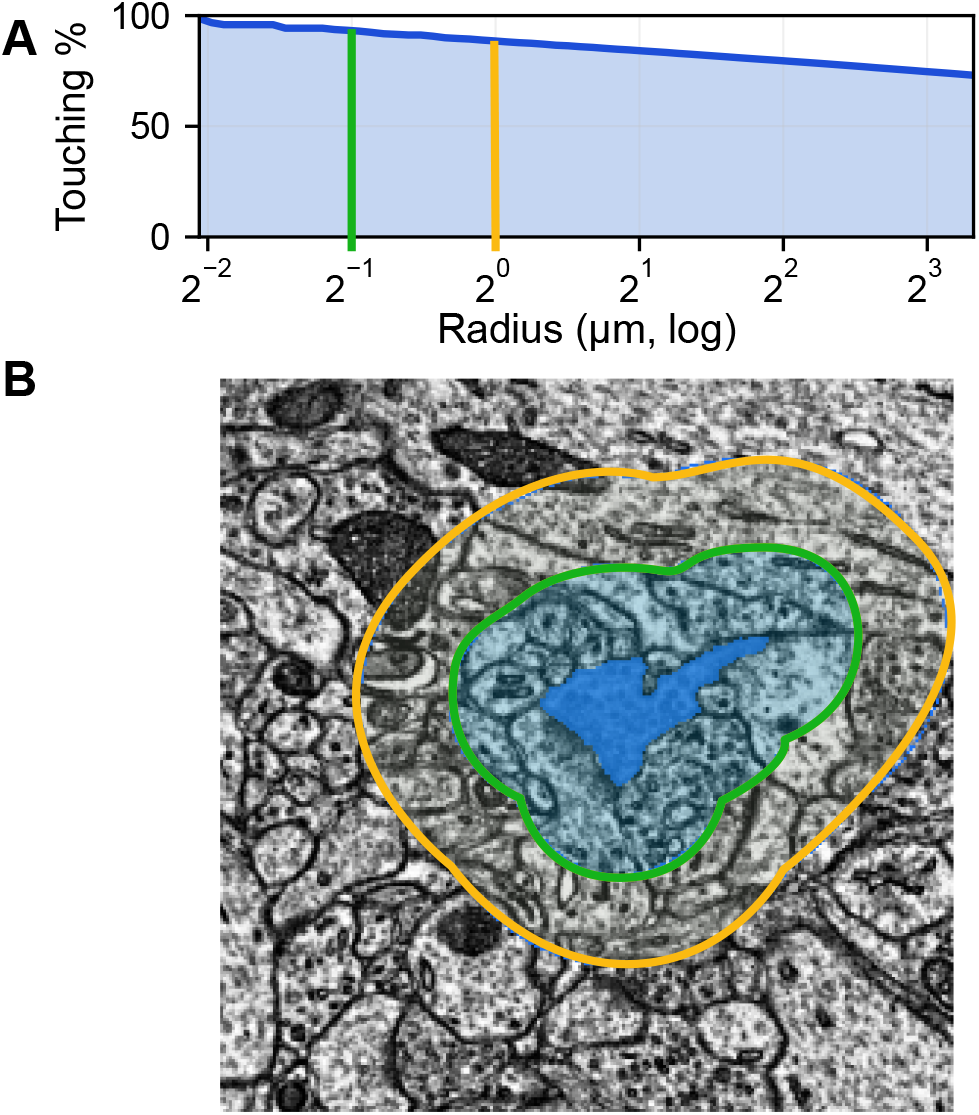
Most nearby cells are already direct contacts. **A**. Fraction of nearby typed pairs that are already direct contacts, as a function of radius. **B**. Representative EM slice showing one neuronal segment (blue) and its 0.5 and 1 *µ*m halo shells (light blue).

### Astrocytes form a cortex-spanning syncytium that contacts the majority of neurons

Prior work has hypothesized an astrocytic syncytium, a large connected network of astrocytes linked by gap junctions.^23,24^ We asked whether the cortical contactome contains evidence for such a structure, and whether it has the necessary scale to influence the neuronal population. We found one dominant astrocyte network that spans the full volume (**Fig. 4**) and directly contacts most typed neurons. Astrocyte organization therefore does indeed operate at tissue scale.

**Figure 4.**
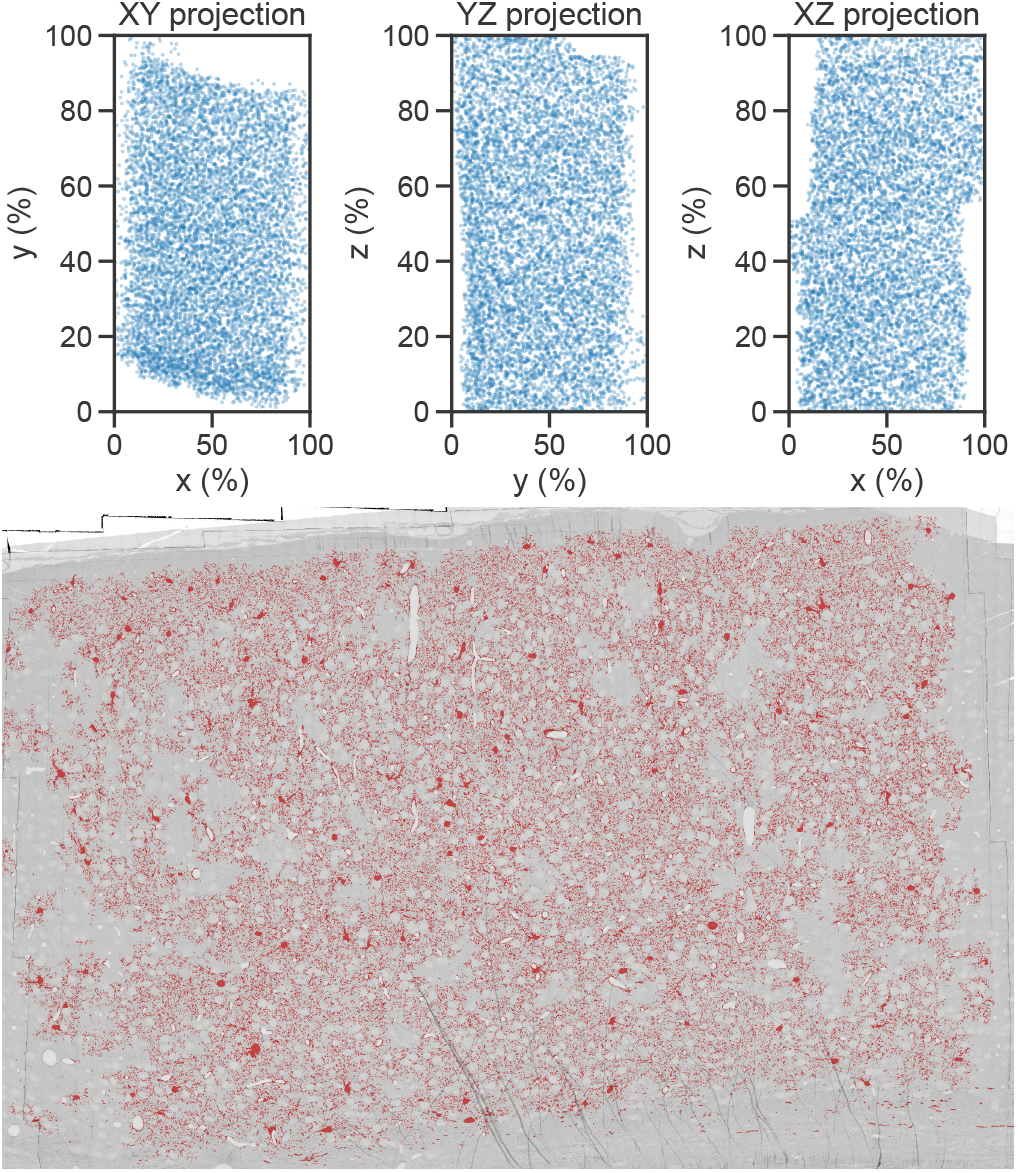
Syncytial astrocytes span the cortical volume. Top: Syncytial astrocyte positions projected into the XY, YZ, and XZ planes, illustrating the uniform distribution of the system across the volume. Bottom: Central microscopy slice with syncytium segmentation overlaid in red. Syncytial astrocytes are distributed throughout the volume rather than confined to a small region, and their processes are interleaved throughout tissue.

This was not a loose collection of local clusters. Among 7,562 total astrocytes, the largest connected component contained 6,146 cells, while the remaining components were much smaller and included 1,184 single-tons (in general as a result of proofreading errors and edge-effects at the volume boundary). This pattern is consistent with a dominant astrocytic syncytium-like network.^23^ This network was also spatially ubiquitous as a tissue-scale structure: Syncytial astrocytes are distributed across the cortical volume (**Fig. 4**), and a central EM slice shows astrocytic processes interleaved throughout the surrounding neuropil rather than confined to isolated patches (**Fig. 1B**).

We next asked how much of the neuronal population this network could actually influence directly. We found that the dominant component directly contacted 61,825 of 71,993 neurons (85.88%), leaving only 10,168 (14.12%) not directly apposed, likely due to annotation- and proofreading edge-effects (*Supplementary Table 3*).

### Synapses exist in a glial microenvironment

Prior work has shown that glia are often part of the synaptic apparatus rather than mere bystanders.^10–12^ However, the extent of synapses that have glial processes nearby remains unanswered. We therefore asked whether synapses in cortex usually have nearby glial processes, and how common glial participants are in the local synaptic microenvironment. We found that glial observers were quite common near synapses and became nearly universal within single-micron distances (**Fig. 5**). Synapses therefore sit almost ubiquitously inside a glial microenvironment, not in isolation.

**Figure 5.**
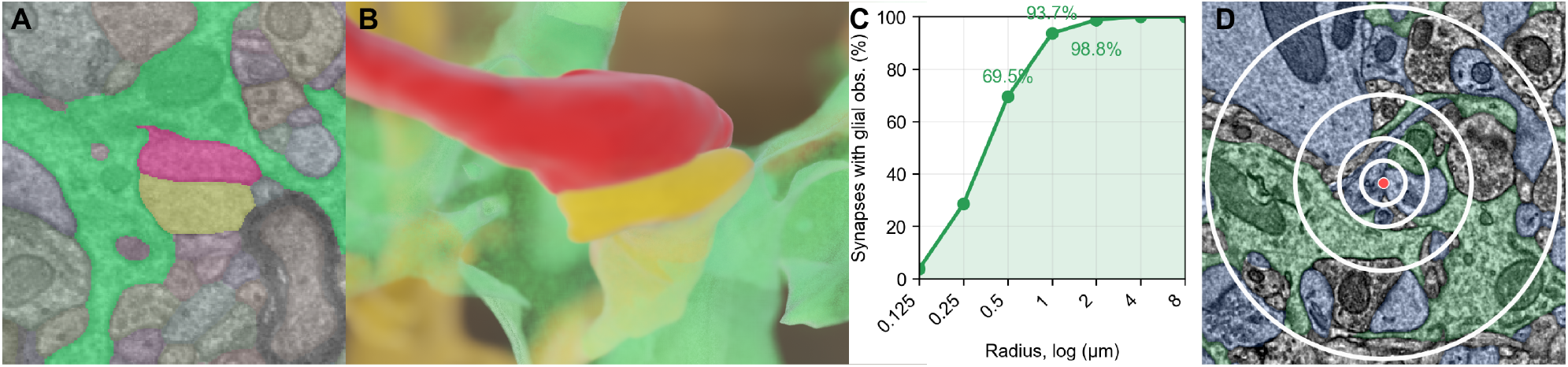
Glial observers are near-universal at cortical synapses. **A**. Electron micrograph of a synapse (red, yellow) with nearby glial processes (green). **B**. 3D reconstruction of the same synapse and nearby glia. The synapse is situated in a glia-heavy microenvironment. **C**. Fraction of synapses with at least one glia nearby, as a function of radius. **D**. EM slice centered on one synapse. Blue = neuronal participants, green = glial segmentation. Rings mark 0.25, 0.5, 1, and 2 µm radii.

To quantify how common this effect is, we sampled *n* = 1, 000 synapses from the interior of the cortical volume and counted third-party “observer” cells within increasing radii around each synapse (excluding the pre- and postsynaptic partners). We found that 93.70% of sampled synapses had at least one glial observer within 1 *µ*m, and 98.80% within 2 *µ*m (**Fig. 5C**). Thus, close glial apposition was the rule rather than the exception.

We next asked whether glia were merely ambient “background” support or meaningful participants in the local environment around synapses. We found that neurons dominated the immediate vicinity of synapses by absolute count, but glia occupied more of the nearby surrounding volume than their cell counts alone would suggest. In other words, glia were enriched immediately surrounding the synaptic site, relative to their ambient density in the tissue. At 1 *µ*m, glia accounted for 16.32% of observer cell counts but 23.19% of the volume. And nearly all synapses (93.70%) had glial participation within 1 *µ*m. Thus, the immediate synaptic neighborhood is highly glia-enriched in nearly all cases.

## Discussion

Synaptic connectomes are powerful, but they capture only one abstraction of cellular relationships in the brain. Here we used the mouse visual cortex contactome to study a broader field of physically available interactions. Neurons contact 5 − 10× more neurons than they synapse with, astrocytes form a dense network, and synapses almost always have glia within 2 µm. Together, these results reveal relational structure that is invisible in the synaptic graph.

This extra structure is not just a denser version of the connectome. Most nearby cells are already realized as direct contacts, but only a subset of contacting pairs are synaptically connected. Contact therefore appears to define a field of physically available interactions from which synapses are selected, consistent with a Peter’s-rule-like view in which adjacency provides the substrate and synapses occupy only a low filling fraction of the contact graph.^13^ The contactome may preserve the local interaction landscape through which developmental synaptic partner selection and synaptogenesis occur.

The same logic extends beyond neuron-neuron pairs. Astrocytes are organized both globally, as one dominant syncytium that touches most neurons, and locally, with glial processes near almost every sampled synapse. Glia appear to be central to the structural context in which cortical communication occurs.

Our manuscript suggests that the synaptic wiring diagram misses much of the relational structure of brains. The connectome tells us who is wired; the contactome tells us who can be in touch.

## Acknowledgements

Research reported in this publication was supported by the National Institute Of Mental Health and National Institute of Neurological Disorders and Stroke under award numbers R24MH114785 and U24NS139927, and by Internal Research & Development funding from the Johns Hopkins Applied Physics Laboratory. The content is solely the responsibility of the authors and does not necessarily represent the official views of the National Institutes of Health.

## Code and Data Availability

All code produced for this study will be made available upon publication.

## Methods

### Datasets

The Layer 2/3 “Pinky100” data volume is a 250 ×140 ×90 *µ*m excerpt of layer 2/3 of the visual cortex of a P36 male mouse.^25–27^ We analyzed fixed public releases of the segmentation and synaptic annotations for this volume from *bossdb*.*org*.^20,28–30^

The 1 mm^3^ MICrONS “Minnie” mouse visual cortex volume is a 1.4 mm × 0.87 mm × 0.84 mm volume spanning visual cortical regions of a P87 mouse.^31^ We used fixed public releases of the volumetric segmentation, synaptic annotations, and cell-type metadata for this volume.^21,28,29^

### Contactome generation

Pairwise cell-cell appositions were extracted from the volumetric segmentations using our generalized volumetric task execution engine *cloudome*^19,32^. Briefly, the volume of interest is divided into chunks, downloaded from BossDB^28^, and processed to account for edge-effects from chunking. Then, 6-way contacts are calculated in units of *nm*^2^ and merged across chunks. Because this workflow operates on local segmentation chunks and connected components rather than materializing the full volume at once, it supports whole-volume contact extraction for arbitrarily large datasets and can be reused across segmented EM volumes with only dataset-specific calibration and metadata changes. Extraction and merge steps were run in distributed batches with out-of-core aggregation with DuckDB^33^. Postprocessing was performed with the *cloudome* database-merge subroutines and results were converted to paired files in Parquet format^34^ (for on-disk compression, parallel reads, and scan operations) and DuckDB (for editability, joins, and other complex queries in SQL). Amortized across our compute jobs, a single 96-core, 1TB RAM node was capable of processing approximately one cubic millimeter per hour. Because source contact-edge weights were reported in dataset-specific voxel-based units, we normalized all pairwise contact areas to *nm*^2^ before downstream analyses using calibration factors derived from voxel geometry: 64×64×40 nm and 128×128×160 nm voxels for the Layer 2/3 and 1 mm^3^ volumes, respectively. All subsequent processing and analyses were performed on a consumer laptop 2023 MacBook Pro M3 Max with 96GB of RAM and 14 cores.

### Graph construction

All contactome graphs were constructed as undirected, unweighted graphs and stored in DuckDB with a *Grand-Graphs* schema for efficient graph queries and joins with cell-type metadata^30,35^.

Graph density is computed in directed graphs as 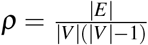 where |*E*| is the edge count and |*V*| is the vertex count of the graph. Density for undirected graphs is defined, 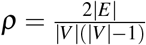.

### Degree distribution

We computed degree distributions directly from pair tables rather than materializing full contactome graphs as in-memory network objects.^30^ The queries used in this analysis are available in the *Supplemental Materials*. For the largest graphs, joins across synapse-contact pairs were executed using batched out-of-core scans.

### Construction of a neuron-neuron subgraph

We composed a mouse visual cortex neuron-neuron sub-graph by restricting the complete contactome to cells with available neuronal type annotations.^36^ Within this table, nearly all synaptic pairs also carried a contact edge, with the remaining exceptions confined to extremely fine-gauge neurites.

### Computing neighborhoods

For the neighborhood analysis, we used the MICrONS Layer 2/3 mouse visual cortex volume. For each typed cell, we defined fixed-distance halos as the sets of other typed cells whose minimum boundary-to-boundary distance fell within thresholds of 0.5, 1, 2, 5, or 10 *µ*m. These distances were estimated on a local coarse segmentation using minimum boundary-voxel-center distances computed with cKDTrees, and the resulting halo neighborhoods were compared with direct contact neighbors to quantify how many nearby cells were already realized as contacts.

